# Skeletal Muscle Histidine Containing Dipeptide Contents are Increased in Freshwater Turtles (*Chrysemys picta bellii*) with Cold-Acclimation

**DOI:** 10.1101/2021.05.31.446418

**Authors:** Eimear Dolan, Daniel E. Warren, Roger C. Harris, Craig Sale, Bruno Gualano, Bryan Saunders

## Abstract

Freshwater turtles found in higher latitudes can experience extreme challenges to acid-base homeostasis while overwintering, due to a combination of cold temperatures along with the potential for environmental hypoxia. Histidine containing dipeptides (HCDs; carnosine, anserine and balenine) may facilitate pH regulation in response to these challenges, through their role as pH buffers. We measured the HCDs content of three tissues (liver, cardiac muscle and skeletal muscle) from the anoxia-tolerant painted turtle (*Chrysemys picta bellii*) acclimated to either 3 or 20°C. HCDs were detected in all tissues, with the highest content shown in the skeletal muscle. Turtles acclimated to 3°C had more HCD in their skeletal muscle than those acclimated to 20°C (carnosine = 20.8±4.5 vs 12.5±5.9 mmol·kg DM^-1^; ES = 1.59 (95%CI: 0.16 – 3.00), P = 0.013). The higher HCD content observed in the skeletal muscle of the cold-acclimated turtles suggests a role in acid-base regulation in response to physiological challenges associated with living in the cold, with the increase possibly related to the temperature sensitivity of carnosine’s dissociation constant and buffering power of the skeletal muscle during anoxic submergence.

**Highlights:**

- pH regulation is a major challenge for overwintering freshwater turtles.
- Histidine containing dipeptides are important intracellular buffers.
- Turtles acclimated to 3°C had higher HCD content than those at 20°C.
- HCDs may be important pH regulators in cold-acclimated turtles.

## INTRODUCTION

The ability to maintain acid-base balance within homeostatic limits is essential for maintaining cellular function (1). Acid-base homeostasis is constantly challenged by various internal and external factors. For example, anaerobic metabolism is required for continued ATP regeneration when turnover exceeds the oxidative capacity of the cell. This results in hydrogen cation (H^+^) accumulation and increased metabolic acidosis (2), which has adverse consequences for numerous cellular processes, including reduced glycolytic enzyme activity, inhibition of oxidative phosphorylation and impaired phosphorylcreatine (PCr) resynthesis (3–5). To avoid these adverse consequences, living organisms have evolved a diverse range of pH regulatory strategies. Intracellular buffers, such as bicarbonate, phosphates, proteins and histidine containing dipeptides (HCDs), provide an important “first line of defence” against intracellular pH perturbations. Simultaneously, “dynamic buffering” is the process by which excess H^+^ is removed from the cell via Na/H^+^ exchangers and monocarboxylate transporters (6, 7).

In vertebrates, the HCDs carnosine (beta-alanyl-L-histidine), along with its methylated analogues balenine (beta-alanyl-1-methyl-L-histidine) and anserine (beta-alanyl-3-methyl-L-histidine), are important intracellular buffers with acid dissociation constants (pKa’s) at physiological temperatures (*i*.*e*., 36°C) that render them ideally placed to buffer across physiological pH ranges (8) - for skeletal muscle this is approximately 7.1 – 6.5 (9, 10). Previous studies indicate that these dipeptides are abundant in the skeletal muscle of species with a large capacity for anaerobic energy metabolism, and that have adapted to tolerate high acid loads (11–13). These species include sprinters, such as thoroughbred racehorses and greyhounds (14); avian species with a limited ability for aerobically fueled flight, such as chickens (15, 16); and aquatic mammals that undergo prolonged periods of hypoxia while diving, such as blue or fin whales (17).

North American pond turtles have a remarkable tolerance to hypoxia (to the point of anoxia), during which they experience major challenges to pH homeostasis (18). As such, they represent a fascinating model to investigate pH buffering. During winter, many of these turtles, especially those found in higher latitudes, are forced to overwinter in anoxic water at the bottom of small ponds and swamps. The western painted turtle (*Chrysemys picta bellii)* can survive anoxia at 3°C for more than 170 days, despite oxygen levels being undetectable (18, 19). This remarkable ability to withstand anoxia results from three main evolutionary adaptations: extreme metabolic suppression, large tissue glycogen stores, and a marked capacity to withstand metabolic acidosis (20, 21). Painted turtles can tolerate very high circulating lactate, with plasma levels of up to 200 mM recorded (21), while blood pH falls to ∼7.2 from normal pH of 8.1 at 3°C (18), representing a remarkable capacity for buffering and pH regulation. To put this into context, humans undertaking exhaustive exercise experience plasma lactate increases of ∼14-18 mM, concomitant to a large export of H^+^ out of the muscle, which generally leads to a reduction in pH from approximately 7.4 to 7.1. Turtles’ buffering ability is largely achieved via the shell (22), which buffers pH by releasing calcium and magnesium carbonates, and via the direct uptake of lactate and H^+^ by the shell (23). Less certain in these species is the contribution of intracellular physicochemical buffers in unmineralized tissues, such as the HCDs, which have previously been reported to be abundant in species who experience large challenges to acid-base regulation (13).

In addition to the challenges that extreme hypoxia poses, these ectothermic vertebrates also hibernate in near freezing conditions, which also has implications for pH regulation, particularly with regards to the charge state of histidyl residues within proteins. Reeve’s alpha-stat hypothesis states that ectotherms shift intracellular and extracellular pH according to temperature in order to maintain constant imidazole ionization, also called alpha (24, 25). HCDs comprise the majority of these intracellular imidazole compounds (11) and as such are likely to contribute toward maintenance of alpha. Therefore, painted turtles are exposed to two major acid-base stressors while hibernating during winter – anoxia and low temperatures. Theoretically HCD content may contribute toward defending against both of these stressors, but little is currently known about the HCD content of these ectothermic vertebrates, nor whether these contents are affected by temperature. The aim of this exploratory study, therefore, was to determine the skeletal muscle, liver and heart HCD content of freshwater western painted turtles who were acclimated to either 3°C or 20°C.

### Animals

Ten adult painted turtles (*Chrysemys picta bellii*; Niles Biological, Sacramento, CA, USA) of both sexes were acclimated to either 3°C (n = 5) or 20° C (n = 5). Prior to temperature acclimation they were maintained for 1-3 months in large tanks filled with partially dechlorinated St. Louis municipal tap water under natural Minnesota photoperiod. The turtles had access to a drying platform, an incandescent light bulb for basking, and a 10W UVB light for nutritional purposes. Air and water temperatures were maintained between 18-22°C. During this time, turtles were fed commercial turtle pellets *ad libitum* three times per week. For the temperature acclimation, the 20°C turtles (n = 5) were transferred to an ∼80-liter temperature-controlled aquarium with water temperature thermostatted to 20°C (YSI Model 72) and were held there, without food, for 36-48 hours. The 3°C turtles (n=5) were placed into an ∼80-liter aquarium with water temperature initially thermostatted to 20°C. The set temperature was lowered by 2°C per day for just over 8 days until it reached 3°C, and then held there for an additional two weeks. The turtles were not fed during the acclimation period. All procedures were approved by the Saint Louis University Institutional Animal Care and Use Committee (IACUC protocol 2198).

### Tissue sampling and preparation

Turtles were euthanized by rapid decapitation. After the plastron was removed with a bone saw, samples of ventricle, liver and pectoralis muscle were removed and immediately flash-frozen with freeze clamps pre-chilled in liquid nitrogen, and then stored at −80°C until analysed. All samples were subsequently lyophilised and powdered before extraction was performed using perchloric acid [HClO_4_], EDTA and potassium bicarbonate [KHCO_3_] (26). The neutralised supernatant was collected using a centrifugal filter (0.2 µm), checked to ensure a pH close to 7 and then stored at −80°C until high-pressure liquid chromatographic (HPLC) analysis.

### Chromatographic determination of histidine-containing dipeptides

The HCD content was determined by HPLC (Hitachi, Hitachi Ltd., Tokyo, Japan), as per Mora et al. (27) using an Atlantis HILIC silica column (4.6×150 mm, 3 μm; (Waters, Massachusetts, USA). All chromatography was conducted at room temperature. Samples were analysed in duplicate and injected via an auto sampler using a loop injection method. Two mobile phases were used. Mobile phase A: 0.65 mM ammonium acetate, in water/acetonitrile (25:75) (v/v). Mobile phase B: 4.55 mM ammonium acetate, in water/acetonitrile (70:30). The pH of both solutions was adjusted to 5.5 using hydrochloric acid and thereafter filtered under vacuum through a 0.2 μm filter membrane. The separation condition comprised a linear gradient from 0 to 100% of solvent B in 13 min at a flow rate of 1.4 mL·min^-1^. Separation was monitored using an ultraviolet detector at a wavelength of 214 nm. The area under the curve (AUC) for carnosine was obtained and used to estimate the content by comparison to standards of 100, 250, 500 and 1000 µM. The in-house variability of the extraction and analysis methods is 4.0 and 2.5% (28). Another peak in close proximity to carnosine was detected in several samples, which was likely to be balenine and/or anserine (27). To determine which of these it was, several samples were spiked with anserine, and the subsequent retention times showed anserine not to be the unidentified peak (retention times, carnosine: 8.5 min; unidentified peak: 9 min; anserine: 9.5 min), meaning that it was balenine (27). Balenine standards were unavailable so quantification could not be performed. Instead, balenine is reported as AUC and relative to carnosine AUC.

### Data analysis

Data were analysed using the SAS statistical package (SAS® University Edition, SAS Institute Inc., USA), and are presented as mean±1SD. HCD content (total HCD AUC and carnosine content mmol·kg DM^-1^) were analysed using mixed model analysis with animals assumed as a random factor and tissue (3 levels; liver/ventricle/*m. pectoralis*) and environment (2 levels; 3 or 20°C) assumed as fixed factors. Tukey–Kramer adjustments were performed when a significant F value was obtained. Results were interpreted according to the statistical probabilities of rejecting the null hypothesis (H0) and in the following categories: p > 0.1: no evidence against H0; 0.05 < p < 0.1: weak evidence against H0; 0.01 < p < 0.05: some evidence against H0; 0.001 < p < 0.01: strong evidence against H0; < p < 0.001: very strong evidence against H0 (29). Effect sizes (ES) were calculated as the mean difference between the two groups of turtles, divided by the pooled standard deviation and are reported alongside their 95% confidence interval. The theoretical buffering contribution of the observed HCD content for a reduction of 0.6 pH units was calculated using a derivation of the Henderson Hasselbalch equation (14) namely: β_HCD_ ={[HCD]/(1 + 10^^(pHi–pKa)^)} – {[HCD]/(1 + 10^^(pHi–pKa)^)}. For this calculation we assumed the physiologically relevant pHi range was 7.1 down to 6.5 at 20°C (30) and 7.4 to 6.8 at 3° (31) and pKa’s for carnosine of 6.702 and 7.209 at 20° and 3°C (32).

## RESULTS

### HCD Content

HCDs were detected in all examined tissues (Figure 1, Panel A). Carnosine was found in the *m. pectoralis* (16.09 ± 7.00 mmol·kgDM^-1^) and balenine in the liver, whilst both balenine and carnosine (6.08 ± 2.95 mmol·kgDM^-1^) were found in the ventricle (contents reported as the mean of all animals). Visual inspection of the chromatograms suggested that very small amounts of balenine were present in two of the *m. pectoralis* samples and very small amounts of carnosine in two of the liver samples, although these were below the limits of interpolation and quantification of the detection software. HCDs were more abundant in *m. pectoralis* than in either of the other two tissues (p ± ≤ 0.001), which did not differ in their total HCD content (p = 0.813; Figure 1B). There was weak evidence that temperature influenced the total HCD content (expressed as combined AUC) combining all tissues (p = 0.092), and there was some evidence of a tissue by temperature interaction (p = 0.021). Post-hoc analysis revealed no differences between temperature conditions for liver (p = 1.00) or ventricle (p = 0.99), but there was some evidence that the turtles acclimated to 3°C had a higher total *m. pectoralis* HCD content than those kept at 20°C (p = 0.023). Turtles maintained at 3°C had a carnosine content of 20.8 ± 4.5 mmol·kg DM^-1^, compared to a content of 12.5 ± 5.9 mmol·kg DM^-1^ in those maintained at 20°C (ES=1.59 (95% CI: 0.16 – 3.00), p=0.013; Figure 1, Panel B). This equated to a buffering contribution of 6.83 ± 1.47 and 4.10 ± 1.93 mmol·kg DM^-1^·(0.6 pH unit)^-1^ for 3 (pHi: 7.4 – 6.8) and 20°C (pHi: 7.1 – 6.5; p = 0.04; ES: 1.59).

**Figure 1:**
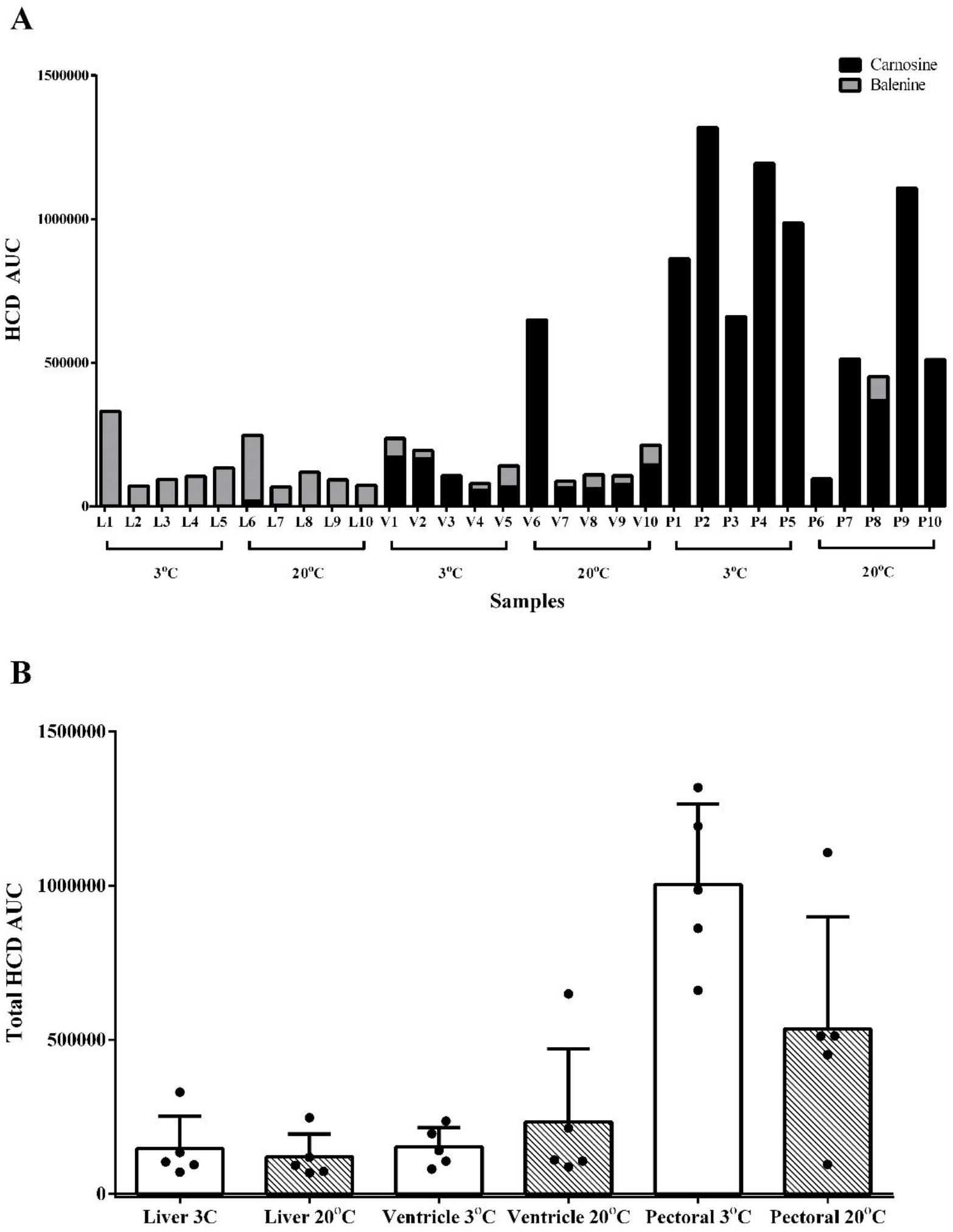
**Panel A:** Individual area under the curve (AUC) HCD content of the turtles maintained at 3 and 20°C. L = liver; V = ventricle and P = Pectoralis. **Panel B:** Mean area under the curve (AUC) HCD content of turtles maintained at 3 or 20°C.

## DISCUSSION

The purpose of this study was to determine the intracellular HCD content of anoxia-tolerant painted turtles acclimated to either 3 or 20°C. The turtles acclimated to 3°C had a higher *m*.*pectoralis* carnosine content than those maintained at 20°C, while liver and cardiac HCD contents were not different. This indicates that intramuscular carnosine content may be instrumental in the adaptive response of these ectotherms to a cold environment.

HCD content varies widely between species and is abundant in the skeletal muscle of those with highly evolved capacities to withstand exercise-induced or environmental hypoxia (11, 13). As such, it might have been expected to observe very high HCD levels in painted turtles, given their remarkable capacity to withstand extreme hypoxia (to the point of anoxia), during which they accumulate blood lactate to levels approaching 200 mM (21). This was not, however, the case and the painted turtles investigated herein had low HCD contents when considered within the context of other species (see figure 2). Even compared with other reptiles, painted turtles have unremarkable HCD contents, with comparable levels to those reported in green sea turtles (*Chelonia mydas;* Suborder Cryptodira) and eastern long-necked turtles (*Chelodina longicollis;* Suborder Pleurodira) (33, 34). The reason for this relatively low intramuscular carnosine content, despite the extreme challenges to acid-base regulation that these animals face, is unclear, but it could relate to the large availability of other mineralized buffers, along with the length of time across which acidosis occurs in this species. Turtles, and painted turtles in particular, can use the calcium and magnesium carbonate stored in their mineralized tissues (*i*.*e*., shell and skeleton) to buffer the acidosis that slowly accumulates over months (23). In contrast, the acidosis incurred by the sprint or diving animals previously reported to be abundant in HCDs (12, 13) is more acute, and occurs when intracellular H^+^ generation is in excess of that which can be actively transported out of the cell. As such, intracellular buffering agents, including tissue HCD contents, may have a greater physiological importance for these animals as opposed to turtles, who experience a more gradual and prolonged acid base stressor while hibernating. In support of this assertion are data suggesting relatively low non-bicarbonate buffering capacities in turtles (35) compared to cetaceans, such as whales (36), who experience the dual challenges of locomotion and hypoxia while diving. In contrast, turtles remain largely stationary when hibernating in anoxic conditions and appear to have solved the buffering problem by exporting H^+^ to the circulation, where it is subsequently buffered by calcium and magnesium carbonates released by the shell, or taken up directly (as lactate and H^+^) and buffered by the shell (23).

**Figure 2:**
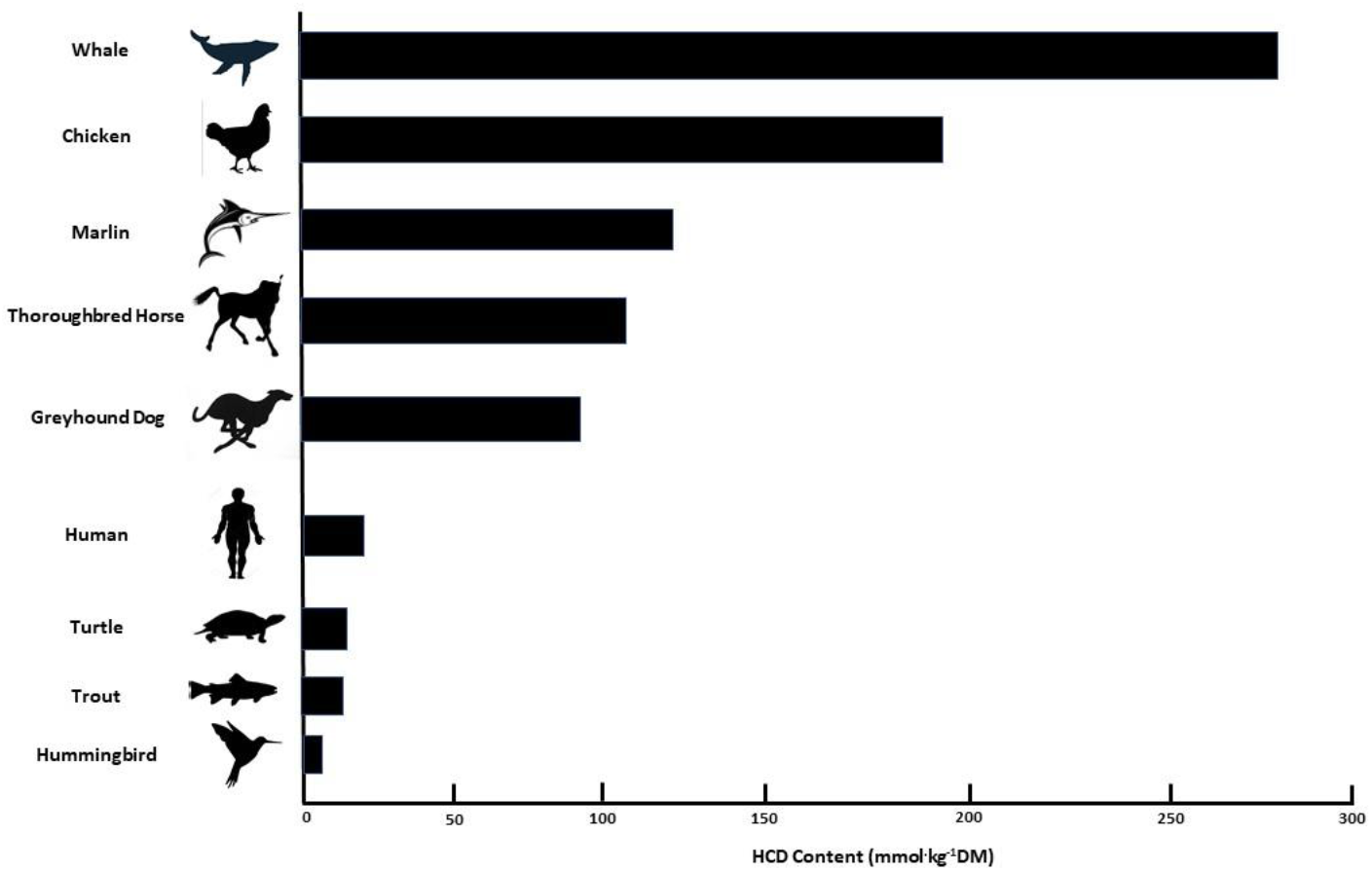
Skeletal muscle HCD content of various species *<u>Note:</u>* The contents shown are indicative, and likely to vary based upon species sub-type and on the muscle type. More detailed overviews of HCD variation in different species are provided elsewhere (11–13).

The higher skeletal muscle carnosine content observed in the cold-acclimated turtles implies a regulatory role for the HCDs in adapting to this stressor. Skeletal muscle forms the largest mass of unmineralized tissue in the turtle, so higher muscle carnosine content could have quantitatively important consequences for acid-base regulation, assuming that it is the predominant histidylated peptide contained therein. Turtles have long been known to be alpha-stat regulators, which means they decrease their PCO_2_ in order to increase their relative alkalinity for the purpose of defending the dissociation fraction (alpha) of the imidazole functional groups in histidine residues within their proteome, thereby preserving their charge and conformational states (24). Based on Reeves (24), the the small change in MCarn observed in the present study would have minimal impacts on intracellular pH or Pco_2_. Thus, the most important effect will be on the buffering power of the muscle. The pKa of carnosine is increased in colder conditions (7.209 at 3°C vs 6.702 at 20°C) which would contribute to the maintenance of the increased alkalinity, given that each temperature-specific pKa is within the mid-range of the assumed pHi at 3°C (7.4 – 6.8) and 20°C (7.1 – 6.5°C). Estimation of the theoretical buffering contribution of the observed MCarn content indicated a higher buffering capacity in the cold-acclimated turtles, again suggesting that this dipeptide may play an important role in acid-base regulation at this temperature. Increases of the magnitude observed herein are roughly comparable to those observed in humans in response to commonly used BA dosing protocols (38), and would necessitate either a large increase in carnosine synthesis, a large reduction in carnosine degradation, or perhaps a combination of both (37). Given that no external BA source was available to these turtles, increased synthesis could only have occurred via increased endogenous production, alongside an increase in carnosine synthase activity. This seems unlikely considering the time-periods that were investigated, and a reduced degradation rate may have contributed (at least in part) to the observed increases. This is, of course, speculative and further research is required both to confirm our findings, and to explore the biokinetics of cold-induced MCarn increases in these ectothermic vertebrates. It is also important to highlight that the numbers of turtles included in this study was small (n = 5 per group), which does increase the risk of sampling error. As such, caution must be applied when interpretating the reported magnitude of increase.

Although the importance of the HCDs to intracellular acid-base regulation is well-recognised, these dipeptides are also thought to contribute toward a number of other biological processes that may be relevant to the dual challenges of anoxia and cold temperatures. Carnosine may play a role in metal ion chelation, antioxidant activity, protein carbonylation and glycoxidation, nonpolysomal proteolysis, and nitric oxide metabolism (18). Of these, carnosine’s antioxidant activity may be most relevant to severely hypoxic and anoxic tissues like those in overwintering turtles, which, theoretically, experience an increase in ROS from xanthine oxidase activity with the reperfusion of oxygen during spring emergence (39). The higher carnosine content, as observed in the cold-acclimated turtles, could, theoretically, increase the antioxidant activity of skeletal muscle and reduce any ROS-mediated injury that might occur during tissue reperfusion.

An interesting finding was the distinct HCD profile within the different tissues. It was unsurprising that skeletal muscle had the largest HCD content, as it has previously been estimated that approximately 99% of the total carnosine is located in this tissue (11). In studies of this kind, three HCD forms are commonly detected in different species, namely carnosine, and its methylated analogues balenine and anserine. Most species contain at least two HCD forms and this varies largely between phylogenetically distinct species, but is similar within the same, or congeneric, species. Only carnosine and balenine were identified in painted turtles, with the latter being absent in skeletal muscle. This is similar to green sea turtles and leatherback turtles (both Cryptodira) (33), but different from eastern long-necked turtles (Pleurodira) (34). The uniformity of HCD distribution of different types in different tissues implies that they may exert type- or tissue-specific effects. Herein, turtles had small amounts of balenine in the liver (AUC), greater amounts of carnosine in the skeletal muscle and a mixture of both balenine and carnosine in the ventricle. The significance of this variation is uncertain, but some biological differences have been reported between the HCD analogues. For example, each HCD has a subtly different pKa (estimated to be 7.03, 7.04 and 6.83 for anserine, balenine and carnosine at mammalian temperatures, *i*.*e*., ∼37°C), while distinct anti-radical effects of the different HCD analogues have also been suggested (40, 41). The functional relevance of these subtle physiological differences is unknown and represents an interesting avenue for future research.

In conclusion, we measured HCDs in the ventricle, liver and *m*.*pectoralis* of the freshwater turtle, with balenine predominating in the liver, carnosine in the skeletal muscle and a mixture of the two in cardiac tissue. Tissue HCD content was relatively low compared to other species with large buffering requirements, implying that overall intracellular buffering may not make a substantial contribution to these turtles’ remarkable capacity to withstand oxygen deprivation. Nonetheless, the HCD content of the *m. pectoralis* was higher in those turtles that were acclimated to 3°C, as opposed to 20°C, which implies a regulatory role for the HCDs in responding to challenges to acid-base homeostasis that occur at colder temperature.

## Declarations

The is work was partially funded by a National Science Foundation CAREER grant (1253939) awarded to Daniel Warren. Eimear Dolan (2019/05616-6 and 2019/26899-6), Bryan Saunders (2016/50438-0) and Bruno Gualano (2017/13552-2) are financially supported by the *Fundação de Ampara a Pesquisa do Estado do São Paulo* (FAPESP). Bryan Saunders has received a grant from Faculdade de Medicina da Universidade de São Paulo (2020.1.362.5.2). None of the authors have any conflict of interest to declare.

